# Brain-wide Organization of Post-Synaptic Sites: Three Principles

**DOI:** 10.64898/2025.12.31.697246

**Authors:** Yina Wei, Yuze Liu, Feng Xiong, Fuhui Long, Hanchuan Peng

## Abstract

Synapses are fundamental units of neural computation and structural plasticity in the brain, yet their brain-wide spatial organization in mammals remains largely unknown. Building on our previous mapping and modeling of dendritic spines and axonal boutons, we developed a statistical approach to generate over 17.99 million predicted post-synaptic sites based on the potential arbor contacts (PPSS_PAC_) across the dendritic arbors of 155,743 neurons spanning the entire mouse brain. By analyzing the topological features of these sites, we identified three key organizational principles. First, PPSS_PAC_ exhibits a distinct region-specific spatial pattern at the dendritic branch level, preferentially enriching near the distal ends of dendritic segments, in contrast to the valley-like distribution of pre-synaptic sites along axonal branches. Cross-species validation against independent electron microscopy datasets from mouse and human brains, revealed a conserved and previously unquantified pattern of synaptic organization. Second, the spatial distribution of PPSS_PAC_ effectively captures variations across cell types in the hippocampus, enabling neuron classification based on synaptic architecture. This serves as a complementary mesoscale descriptor to existing classifications based on global neuronal connectivity (source-target relationships) or local dendritic microenvironments. Third, the spatial distribution of PPSS_PAC_, rather than synapse density, shows remarkable correlation with independent whole-brain electrophysiological activity. The spatial organization of PPSS_PAC_ also correlates with several gene expression datasets. These results suggest the post-synaptic pattern could be mediator linking molecular signatures, neural dynamics, and connectivity types of cells. Our study reveals, for the first time, a structural substrate for hierarchical information processing in the brain, providing new insights into neural computation and a foundation for brain digital twin models to support next-generation AI infrastructure.

## Introduction

Synaptic connections are essential for neural communication, providing the structural and functional substrates that support neural networks^1,2,3,4,5,6^. These connections occur at distinct locations along neuronal structures, specifically at the pre-synaptic and post-synaptic sites, where their spatial arrangement critically influences circuit integration and function^7,8,9^. In particular, post-synaptic sites serve as key integration hubs for incoming synaptic inputs^10^, modulating neuronal processing, plasticity, and memory formation^11,12,13^. Their spatial distribution influences how synaptic signals are combined, reflecting functional organization in which inputs related to similar features tend to cluster together^14,15,16^.

The non-random distribution of synaptic sites is fundamental to neuronal function, shaping output through mechanisms such as local filtering, nonlinear integration, and the influence of voltage-gated ion channels^17,18^. Beyond these functional roles, these spatial arrangements may also serve as potential markers for neuronal identity, reflecting the unique characteristics of various cell types. While previous cell classification strategies have relied on global connectivity patterns^4^, or local dendritic microenvironments^5^, we hypothesize that the spatial organization of post-synaptic sites may introduce a complementary mesoscale descriptor. This perspective may also enable the identification of cell types based on the spatial distribution of synaptic inputs along dendrites.

Despite progress in understanding the molecular mechanisms underlying synaptic function and plasticity^19,20,21^, the spatial distribution of post-synaptic sites remains poorly explored. This gap is partly due to the complexity of dendritic structures, which vary across neuronal types and brain regions^22^, as well as the technical challenges associated with large-scale synaptic imaging^23,24^. High-resolution imaging, such as electron microscopy (EM), is required to map synaptic locations accurately. However, the high cost and technical difficulty of scanning large tissue volumes at such resolution, combined with the complexities of subsequent morphological reconstruction, have made large-scale studies of synaptic distribution challenging^25,26^.

Recent advances in synaptic imaging and reconstruction have deepened insights into synaptic organization. For example, specific patterns of excitatory and inhibitory synapses have been identified in layer 2/3 pyramidal neurons^1^, emphasizing local excitatory/inhibitory (E/I) balance. Large-scale initiatives, such as the MICrONS program^27^, have generated dense reconstructions of synaptic sites in the visual cortex, showing the detailed wiring principles of cortical circuits, including structured connectivity patterns among excitatory and inhibitory neurons, high-precision mapping of synaptic partners, and single-neuron connectivity across cortical layers and visual areas. A recent study has provided valuable insights into the relative synaptic density across brain regions^28^, revealing a dynamic and region-specific pattern of synapse density that changes across the mouse life span. Nevertheless, the brain-wide spatial organization of synapses at the single-neuron level remains largely unexplored in mammals.

Although our previous work^2,3^ has mapped brain-wide pre-synaptic distributions, the organization of post-synaptic inputs along dendrites remains unexplored. In this study, we expanded upon this foundational work by introducing a statistical approach to generate a large-scale dataset of over 17.99 million predicted post-synaptic sites (PPSS) based on the potential arbor contacts (PAC) across 155,743 dendrites spanning multiple brain regions. The dataset enables us to analyze the topological features of these predicted synapses and identify the region-specific patterns of synaptic distribution. Furthermore, we validated the synaptic predictions through comparisons with independent multi-modality datasets, including EM-based reconstructions from mouse and human brains. Our study provides critical insights into the structural organization of synapses, advancing our understanding of neural computation and brain function. These insights establish a biological foundation for developing brain digital twins, enabling next-generation artificial intelligence systems.

## Results

### PPSS_PAC_ exhibits distinct spatial distributions within dendritic arbors

The anatomical colocation of axons and dendrites has been widely adopted as a proxy for potential synaptic connectivity^3,29^. Our previous work has proposed a scalable and cross-validated framework to reconstruct single-neuron connectomes and generate the PACs for neuronal connections in mouse brains. Building on this framework, we generated a large-scale dataset of over 17.99 million predicted post-synaptic sites based on the potential arbor contacts (PPSS_PAC_) distributed along dendritic arbors from over 100 mouse brains (**Figure 1A**) to investigate the spatial organization of the synaptic targeting at the whole-brain scale. Specifically, the PPSS_PAC_ dataset was generated using 155,743 morphologically reconstructed dendrites (**Supplementary Table 1**) sampled from multiple brain regions^2^, together with 18,271 axonal reconstructions accessed from publicly available datasets^22,30,31^ (**Methods**).

**Figure 1.**
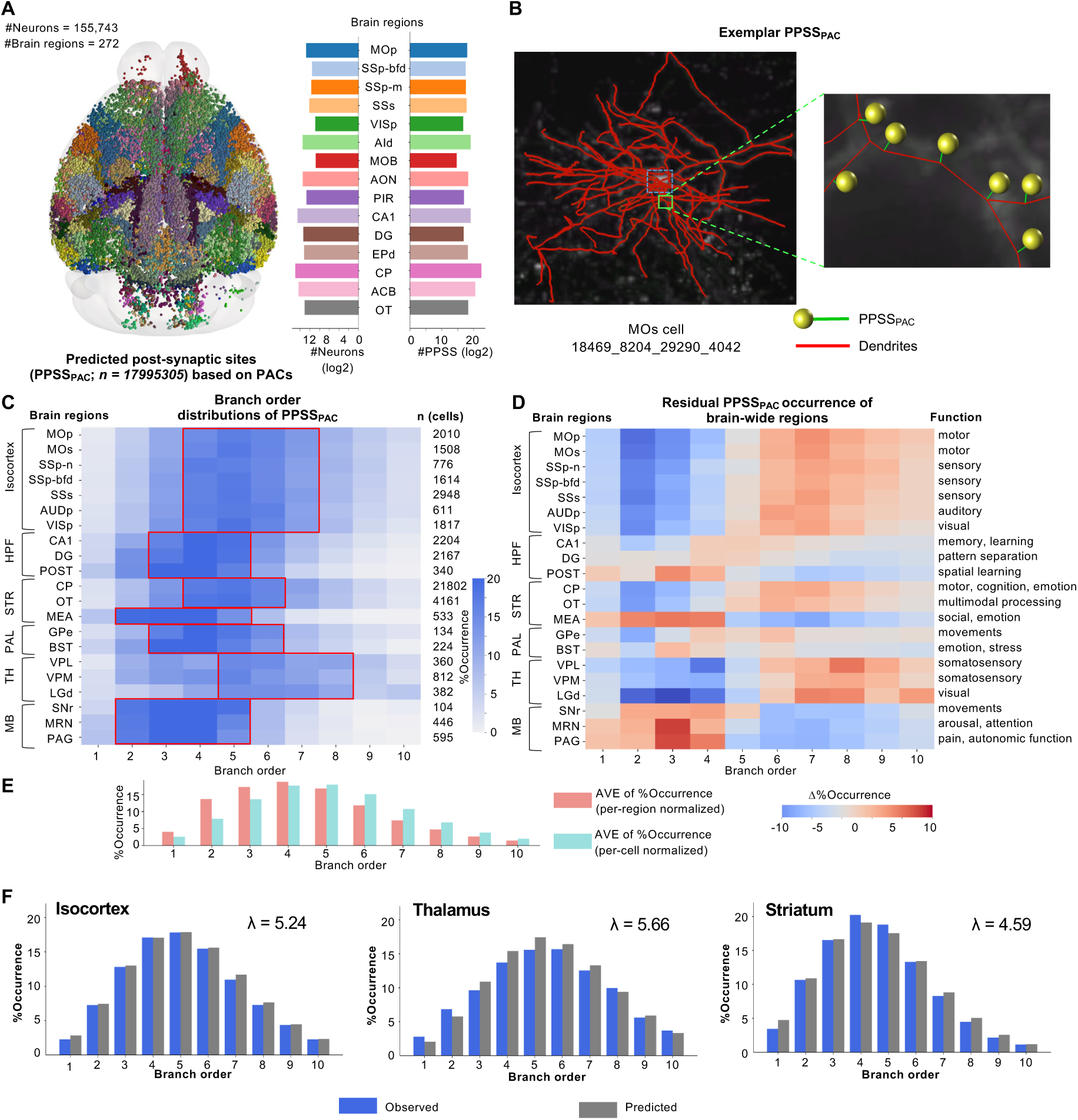
Spatial distribution and branch order patterns of predicted post-synaptic sites (PPSS_PAC_) across 155,743 dendrites from whole mouse brains. **A.** Left: spatial distribution of PPSS_PAC_ in the whole mouse brain. Right: number of reconstructed dendrites and total counts of PPSS_PAC_ identified in main brain regions. **B.** 3D visualization of PPSS_PAC_ along the dendritic branches. Ball-sticks represent the predicted dendritic spines. Red lines represent the dendritic arbors. Blue dotted box highlights the location of the soma. **C.** The distribution of PPSS_PAC_ across dendritic branch orders for different brain regions. The colorbar represents the occurrence of PPSS_PAC_ at each branch order. The red boxes highlight the branch orders with the peak concentration of PPSS_PAC_ for each primary brain region. **D.** The residual PPSS_PAC_ occurrence across dendritic branch orders for brain-wide regions. For each branch order, the values represent the deviation from the mean PPSS_PAC_ occurrence within that branch order. The colorbar indicates the magnitude of these residuals, with positive and negative values reflecting above- and below-mean deviations, respectively. **E.** The histogram represents the distribution of mean PPSS_PAC_ occurrence values across all branch orders. Light red: averaged per region; Light blue: averaged per cell. **F.** Branch order distribution and best-fit Poisson models of PPSS_PAC_ in major brain regions: isocortex, thalamus, and striatum. Colors indicate the real distributions (blue) and best-fit distributions (grey).

To infer the locations of PPSS_PAC_, we first computed the shortest Euclidean distance between each axonal arbor and its nearest dendritic arbor across all neuron pairs. A dendritic compartment was defined as a predicted post-synaptic site (**Figure 1B**) if the distance to its nearest axonal arbor was less than 5μm, a threshold representing the sum of the typical length of dendritic spines and axonal boutons, as reported in previous studies^1,32^.

By applying the spatial proximity-based estimation, we identified more than 17.99 million PPSS_PAC_ and registered them to the Allen Common Coordinate Framework (CCF) version 3 atlas^33^ using the mBrainAligner tool^34^. Through the systematic comparative analysis of synaptic distribution across the whole mouse brain, we observed that the PPSS_PAC_ were broadly distributed in the entire brain, reflecting not only the widespread reach of axonal projections but also the broad spatial distribution of dendritic targets (**Figure 1A**). This widespread distribution of axonal and dendritic arbors enables a comprehensive investigation of the spatial patterns governing PPSS_PAC_ distribution.

Understanding the spatial distribution of synaptic targets along the dendritic arbor is critical for interpreting neuronal integration and circuit function^35,36^. Previous studies of retinal ganglion cells from rabbits and mice have explored the spatial patterns of post-synaptic sites in dendritic structures, showing less specialized distribution^37,38^. Despite these efforts, the spatial organization of post-synaptic sites across diverse brain regions remains largely unexplored^39,40^. To address this, we quantified the average proportions of PPSS_PAC_ occurrences across branch orders in dendrites sampled from brain-wide regions (**Methods**; **Supplementary Figure 1A**; **Supplementary Table 2**).

We observed that PPSS_PAC_ exhibits a distinct, non-uniform distribution along dendritic arbors, characterized by preferential concentrations within specific branch orders rather than a uniform arrangement. For example, peak occurrences of PPSS_PAC_ were found at dendritic branch orders 4–7 in the isocortex, 5–8 in the thalamus, and 2–5 in the midbrain (**Figure 1C**). To account for the sampling bias introduced by the reduced prevalence of higher-order branches, we used an alternative normalization strategy by the number of neurons that possessed branches at that specific order (**Methods**; **Supplementary Figure 1B**). The similar spatial patterns of PPSS_PAC_ observed between the two different normalization approaches (**Supplementary Figure 1**) supported the key observation that the spatial distribution of PPSS_PAC_ is closely coupled to the underlying anatomical layout of dendritic arbors (**Supplementary Figure 1B**). To quantify the spatial distribution of PPSS_PAC_, each dendritic branch was divided into three equal-length segments (**Methods**): proximal (near the branch origins), middle (near the branch centers), and distal (near the branch ends). We observed a pronounced enrichment of PPSS_PAC_ at distal segments (**Supplementary Figure 2**), where synaptic sites preferentially clustered. This distal enrichment may indicate these compartments as functionally critical hubs for receiving and integrating synaptic inputs.

We observed a region-specific organization of PPSS_PAC_, characterized by highly identical branch-order occurrence rates within major regions such as the isocortex (**Figure 1C, red box**). This spatial patterning was consistently observed in residual PPSS_PAC_ (**Figure 1D**) after removal of the global average, whether profiles were averaged per region (**Figure 1E, light red**) or per cell (**Figure 1E, light blue**). These results demonstrate that synaptic wiring adheres to a regionally specific organizational principle, which may underlie specialized computational functions across different brain areas.

To quantitatively characterize this organization, we fitted PPSS_PAC_ from each major brain region with several probabilistic models. The Poisson distribution provided the best fit according to the Bayesian information criterion (**Supplementary Table 3**). We applied Poisson modeling to three major brain structures, including isocortex, thalamus, and striatum, revealing distinct region-specific parameters (**Figure 1F**). Sub-regions analysis further exhibited varied parameter distributions across major regions (**Supplementary Figure 3A**), indicating region-specific synaptic organization patterns. Model validation through moment analysis showed significant inter-regional differences in both expectation and variance (**Supplementary Figures 3B**, **C**, **D**), with the variance-to-expectation ratio consistently near 1, confirming the suitability of the Poisson distribution.

### Spatial proximity and anatomical structure constrain inter-regional PPSS_PAC_ similarity

Our analysis of the region-specific PPSS_PAC_ distributions (**Figure 1C**) revealed the spatial principles underlying how synaptic inputs are positioned along dendritic arbors. To further explore whether different brain regions exhibit distinct patterns of dendritic targeting, we constructed feature vectors for brain regions using the distribution of PPSS_PAC_ occurrences across dendritic branch orders (**Figure 1C-D**; **Methods**). This approach effectively characterized and enabled the comparison of synaptic targeting patterns across brain regions. These feature vectors were then projected into a low-dimensional space using Uniform Manifold Approximation and Projection (UMAP), enabling a comparative overview of inter-regional relationships in synaptic organization.

We observed that brain regions belonging to the same primary anatomical structures, such as the isocortex, thalamus, and striatum, tended to cluster together in the embedding space (**Figure 2A**). This clustering suggests that sub-regions within the same anatomical structures share similar synaptic targeting patterns across dendritic hierarchies, consistent with our initial observations of the region-specific patterns (**Figure 1C)**. These findings provide evidence that synaptic targeting along dendrites is shaped and constrained by anatomical organization, rather than occurring randomly.

**Figure 2.**
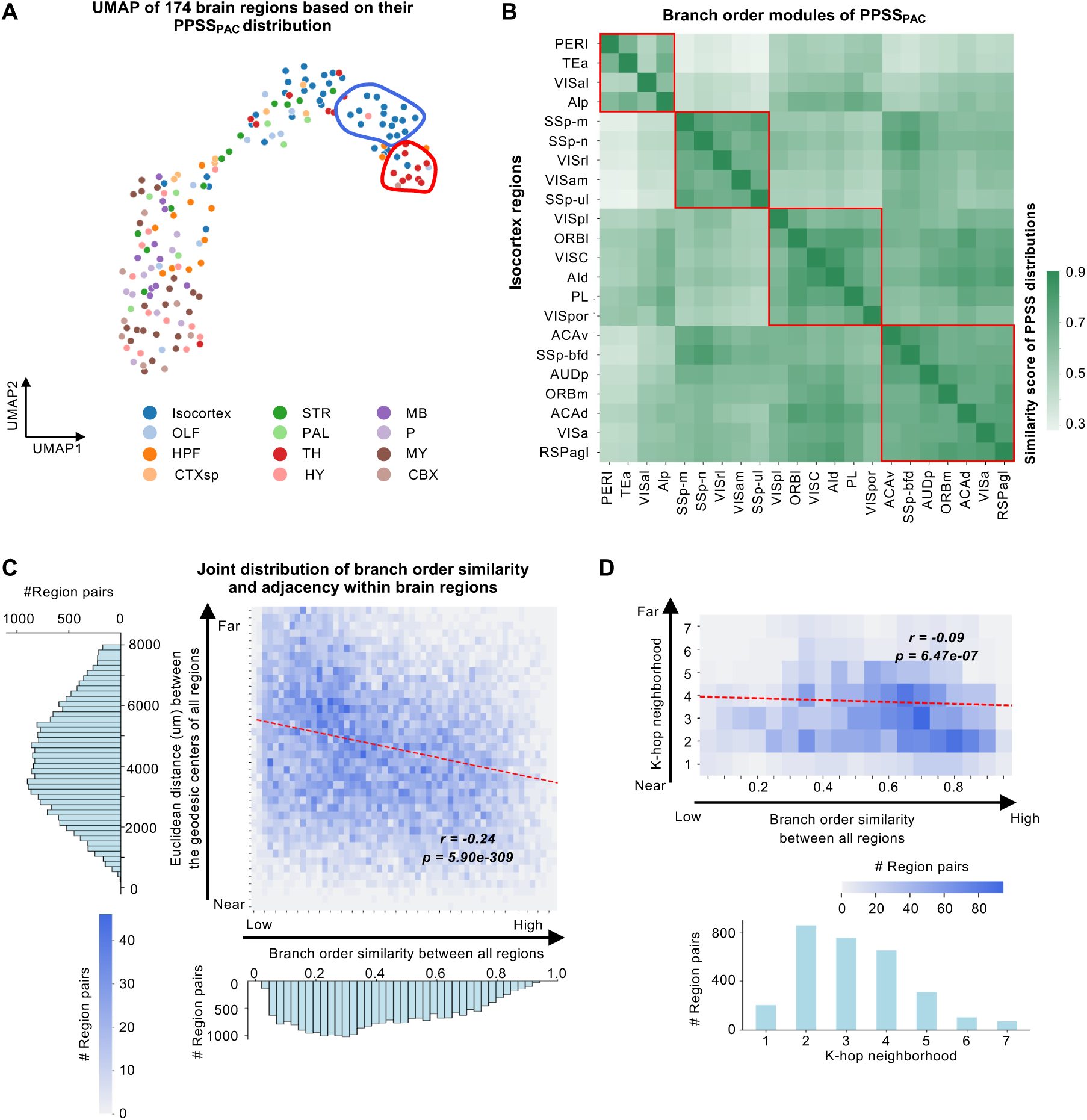
Region-specific similarity of PPSS_PAC_ branch order distributions and their relationship with spatial proximity. **A.** UMAP embedding of the PPSS_PAC_ branch order distributions across 200 brain regions. Each dot represents a brain region, colored according to its primary anatomical structure. **B.** Heatmap showing pairwise similarity of PPSS_PAC_ branch order distributions among Isocortex sub-regions (22 regions shown). Sub-regions are clustered into four modules highlighted by red boxes. **C-D.** Relationship between pairwise similarity in PPSS_PAC_ branch order distributions and inter-regional spatial proximity. Heatmaps show the joint distributions of region pairs based on their PPSS_PAC_ distributional similarity and either spatial distance (**C**, **top right**) or adjacency distance (**D**, **top**). Color bars indicate the number of region pairs at each coordinate. Histograms show the marginal distributions of PPSS_PAC_ distributional similarity (**C**, **bottom**), the Euclidean distances (**C**, **top left**), and regional adjacency (**D**, **bottom**).

Existing studies of cortical neurons showed distinct patterns of synapse distribution, with different types of interneurons targeting specific dendritic domains of pyramidal neurons^41,42^. Therefore, we examined the sub-regions of isocortex in detail to assess whether such synaptic organization extends to finer anatomical scales. We computed the pairwise similarity matrix based on the PPSS_PAC_ branch order features of 43 isocortical sub-regions (**Figure 2B**; **Methods**), resulting in four modules of sub-regions with similar synaptic distributions. It demonstrated that dendritic synaptic targeting follows the specific and non-random spatial patterns within the isocortex, indicating distinct isocortical connectivity patterns characterized by the unique synaptic targeting signatures.

To identify potential anatomical determinants of these inter-regional similarity scores, we evaluated two fundamental spatial relationships within these brain regions (**Figure 2C-D**): (1) physical spatial proximity, measured by the Euclidean distance between region centroids, and (2) direct anatomical adjacency, defined as the minimal number of intermediate regions separating two regions. The two complementary metrics allowed us to assess whether the spatial relationship between two regions affects their synaptic targeting patterns.

We found that inter-regional similarity of PPSS_PAC_ pattern were negatively correlated with both the Euclidean distance (Pearson’s *r* = -0.24, *p* = 5.90×10^-309^) (**Figure 2C**) and adjacency distance (Pearson’s *r* = -0.09, *p* = 6.47×10^-07^) (**Figure 2D**). In other words, brain regions that are closer in 3D space or more directly connected tend to display more similar dendritic synaptic profiles. The strong correlations highlight a distance-dependent organizational principle, linking mesoscale anatomical organization to microscale synaptic arrangement. Together, these findings suggest that spatial, developmental, or functional constraints may act across scales to shape the distribution of synaptic inputs along dendritic trees.

To assess the validity of our PPSS_PAC_ distribution patterns, we also employed ChatGPT-5 to independently analyze brain-wide data and compared its outputs with human-designed analysis. The patterns identified by ChatGPT-5 closely aligned with those from human-designed analysis, further confirming the reliability and robustness of our previous observations (**Supplementary Figure 3A**). Particularly, brain regions were manually categorized into three groups based on the single branch order with the highest PPSS_PAC_ occurrences (**Supplementary Figure 3B, left**; **Supplementary Table 4**). We then provided ChatGPT-5 with the brain-wide PPSS_PAC_ distributions (**Supplementary Table 2**) and a prompt to perform the same grouping (**Supplementary Figure 3B, right**). Accordingly, ChatGPT-5 generated a comparable classification by summing the PPSS_PAC_ occurrences within low-order (branch orders 1-3), middle-order (4-5), and high-order (6-10) ranges for each region, and assigning each region to the category with the highest summed value (**Supplementary Figure 6B, right**). Despite the methodological difference, the resulting classifications were highly consistent. The branch order distribution of brain regions in the UMAP space showed distinct clusters corresponding to the synaptic targeting preferences identified in both the human-driven analysis and ChatGPT’s classification. Brain regions with a preference for PPSS_PAC_ in lower branch orders (1-3) clustered together, while regions favoring middle (4-5) and higher branch orders (6-10) formed separate and distinct clusters. This strong consistency between the spatial patterns derived from manual and ChatGPT-based approaches further underscores the robustness and reliability of our observations.

### PPSS_PAC_ distribution aligns with structural, functional, and vascular features of the brain

In our previous work^3^, we developed two independent approaches to predict single-neuron connections and their associated synaptic contact sites: PPSS_PAC_ based on the potential arbor contact, and PPSS_PB_ derived from predicted boutons (PBs) along their axonal arbors (**Methods**). Here, we implemented these approaches for comparative analysis, generating a dataset of 126892 PPSS_PB_ from the 155,743 dendrites. The two parallel methodologies allowed us to cross-validate these post-synaptic predictions using different structural features of neuronal morphology.

We found that the branch order distributions of PPSS_PB_ displayed specific patterns across brain-wide regions (**Figure 3A**; **Supplementary Table 5**), which closely resembled those observed for PPSS_PAC_ (**Figure 1C**). A quantitative comparison of branch order distributions between PPSS_PAC_ and PPSS_PB_ displayed a strong positive correlation across all regions (Pearson’s *r* = 0.91, *p* = 0.00; **Supplementary Figure 5A**). The agreement between two prediction approaches demonstrated the robustness of our statistical prediction for post-synaptic targeting in whole-brain dendrites.

**Figure 3.**
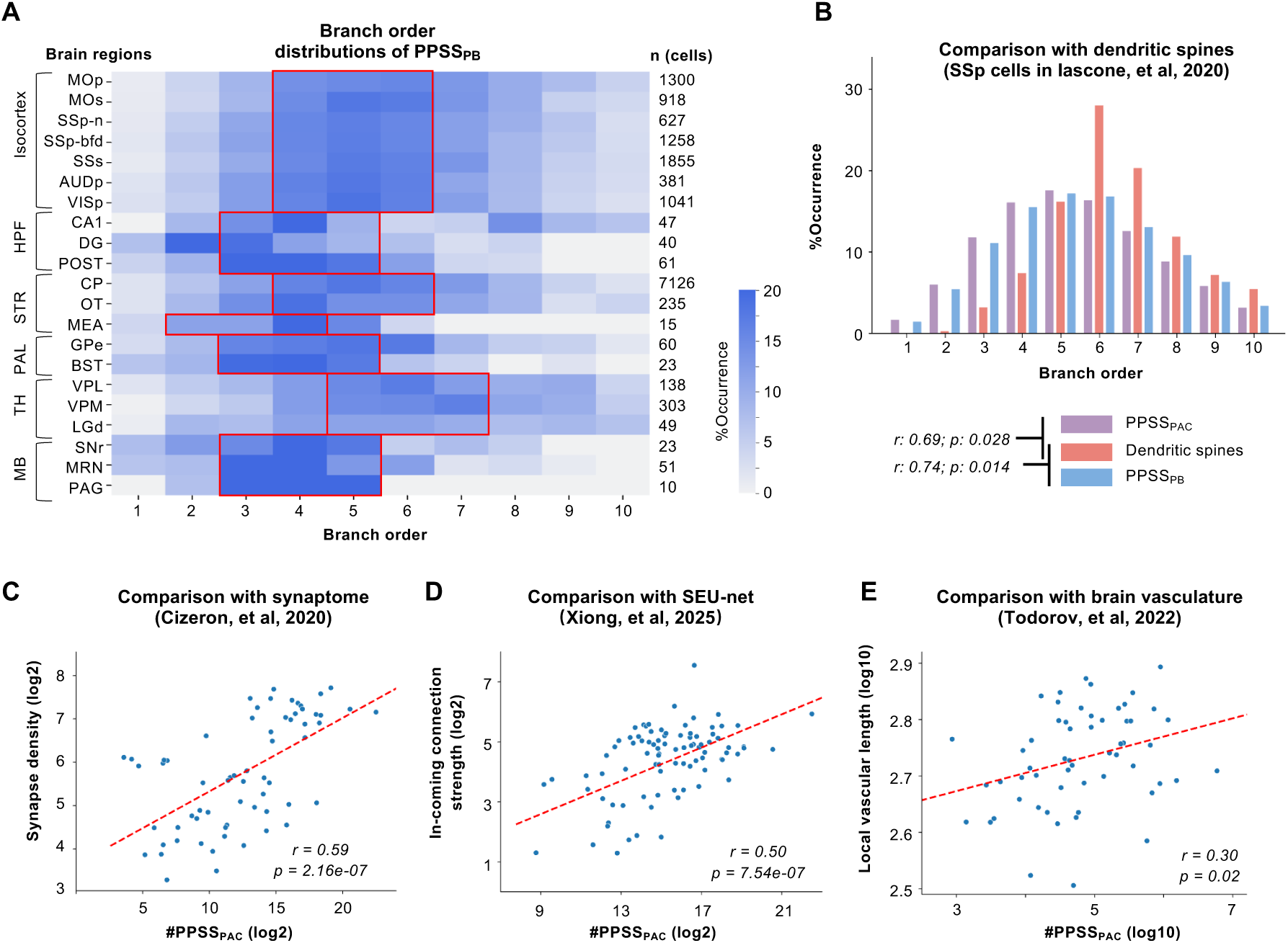
Cross-validation of predicted post-synaptic sites (PPSS_PAC_) across multi-modal datasets. **A.** The distribution of PPSS_PAC_ across dendritic branch orders for different brain regions, where PPSS_PB_ was inferred based on predicted bouton sites on axons. The colorbar represents the occurrence of PPSS_PB_ at each branch order. The red boxes highlight the branch orders with the peak concentration of PPSS_PB_ for each primary brain region. **B.** The comparison of branch order distributions between PPSS and the dendritic spines from SSp neurons. Bar plots show the proportion of synaptic sites across dendritic branch orders. **C**. The correlation between the number of PPSS_PAC_ in each brain region and the relative synapse density obtained from a synaptome atlas. Each dot represents a brain region. **D**. The correlation between the number of PPSS_PAC_ in each brain region and the strength of incoming anatomical connections based on SEU-net. **E.** The correlation between the number of PPSS_PAC_ and the local vascular length in each brain region.

To further validate the spatial distribution of PPSS_PAC_ along dendritic arbors, we analyzed twelve morphologically reconstructed neurons from the primary somatosensory cortex (SSp) with annotated dendritic spines^1^. Quantitative analysis of branch order distributions demonstrated remarkable consistency between predicted and experimentally observed post-synaptic sites (**Figure 3B**). Both PPSS_PAC_ (Pearson’s *r* = 0.69, *p* = 0.028) and PPSS_PB_ (Pearson’s *r* = 0.74, *p* = 0.014) were highly correlated with experimentally measured spine distributions. These results again validated our approach, demonstrating its ability to accurately reconstruct biologically realistic post-synaptic organization across dendritic arbors. We also explored the spine distributions along individual dendritic branches, divided equally into proximal, middle, and distal segments, revealing a characteristic distal enrichment pattern (**Supplementary Figure 5B**). Notably, this biologically observed distribution precisely correlated with the spatial distributions of PPSS_PAC_ mapped to corresponding branches in our reconstructed SSp dendrites.

To evaluate whether the anatomical distribution of PPSS_PAC_ matched an established anatomical distribution of synapses at the whole-brain scale, we compared our PPSS_PAC_ dataset with a whole-brain synaptome atlas^28^, which provided the distribution of relative synapse density in brain-wide regions. Across the 53 brain regions shared by both datasets, we found a positive correlation between regional PPSS_PAC_ counts and experimentally measured synapse density (Pearson’s *r* = 0.63, *p* = 1.55e-10; **Figure 3C**). This correlation indicates that our predictive framework accurately captures key aspects of synaptic organization.

The spatial organization of synaptic inputs along the dendrites is important for the connectivity mappings between individual neurons^43,44^. Therefore, we compared the PPSS_PAC_ distributions with a meso-scale connectome, called SEU-net^3^, which provided a reliable estimation of region-to-region connection strengths. We found that brain regions receiving stronger anatomical inputs also exhibited a greater number of PPSS_PAC_ (**Figure 3D**). This positive correlation (Pearson’s *r* = 0.50, *p* = 7.54e-07) implies that PPSS_PAC_ not only reflects local synapse density but also aligns with the broader principles of the brain-wide connectivity networks.

In addition, we also compared PPSS_PAC_ with the length of blood vessels in each brain region from a mouse brain vascular atlas^45^. It has been hypothesized that vascular distribution is related to brain activity, with brain regions exhibiting stronger activity requiring more blood supply^46,47^. Based on this, we proposed that regions with higher vascular density might also have more synapses. By comparing the vascular length distribution with the corresponding PPSS_PAC_ counts across brain regions, we found a positive correlation (Pearson’s *r* = 0.30, *p* = 0.02), indicating that regions with longer vasculatures may indeed exhibit abundant synapses (**Figure 3E**). This result highlights a potential link between brain activity, vascular support, and synaptic density.

Taken together, these cross-validation analyses, including independent prediction methodologies, reconstructed SSp neurons, brain-wide synapse density distribution, anatomical connectivity datasets, and brain vascular length, not only support the reliability of PPSS_PAC_ predictions but also reveal conserved principles of synaptic organization on the dendritic trees. This multi-modal validation provides strong evidence that PPSS_PAC_ captures fundamental aspects of synaptic organization throughout the brain.

### PPSS_PAC_ distribution captures conserved spatial principles across human and mouse brains

EM provides high-resolution reconstruction of synaptic structures, offering critical benchmark for validating computational models of synaptic connections^48^. To evaluate the biological relevance of PPSS_PAC_, we compared the PPSS_PAC_ with two high-quality EM datasets: (1) the MICrONS dataset^27^, which provides both structural and functional data from the mouse visual cortex (VIS); (2) the H01 dataset^8^, which offers detailed reconstructions of neural circuits in the middle temporal gyrus (MTG) of the human temporal cortex. This cross-species comparisons evaluated whether PPSS_PAC_ recapitulates the spatial distributions of post-synaptic sites across cortical layers and dendritic branch orders observed experimentally.

In the mouse VIS, the PPSS_PAC_ displayed a concentrated distribution within the dendritic branch orders 3-6, with a sharp decline beyond the 6th order (**Figure 4A**). Similarly, the MICrONS dataset showed that experimentally identified synapses were primarily located between branch orders 2 and 5 (**Figure 4B**), with very few synapses observed beyond order 5. The two distributions were significantly correlated (**Figure 4D, left**; Pearson’s *r* = 0.54, *p* = 7.43e-06), supporting the biological plausibility of PPSS_PAC_ in mouse cortex. The subtle differences may be attributable to biological variability, such as age, sex, or genetic background, between the mice used to generate the PPSS_PAC_ and those in the MICrONS dataset, all of which can influence dendritic architecture and synaptic distribution.

**Figure 4.**
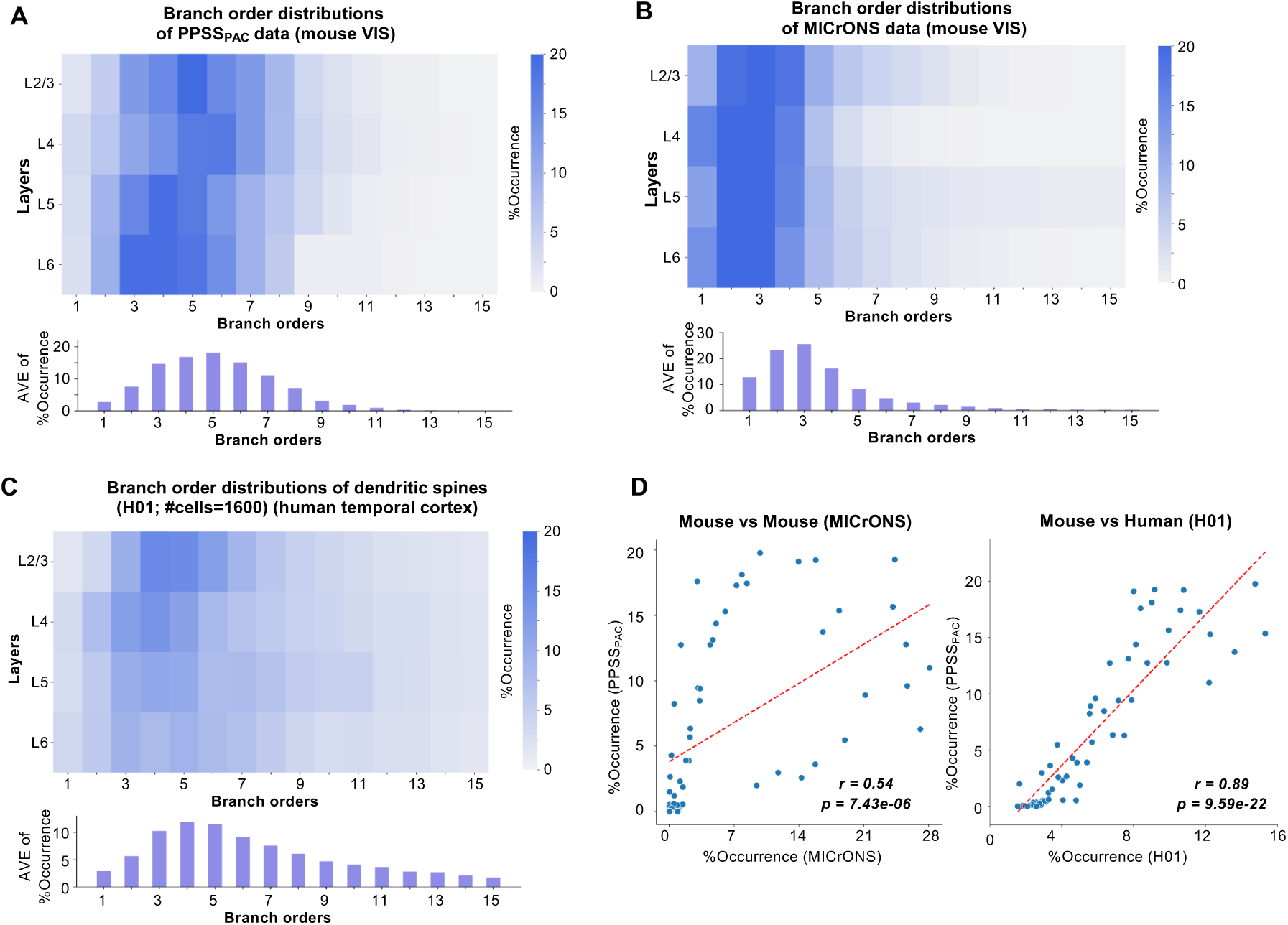
Validating the branch order distributions of PPSS_PAC_ via EM data. **A**. The distribution of PPSS_PAC_ across dendritic branch orders for mouse visual cortex (VIS). The colorbar represents the occurrence of PPSS_PAC_ at each branch order. **B**. The distribution of post-synaptic sites across dendritic branch orders for the MICrONS dataset, reconstructed using EM in the mouse VIS. **C**. The distribution of post-synaptic sites across dendritic branch orders for the H01 dataset, reconstructed using EM in the human temporal cortex. **D**. The comparison between PPSS_PAC_ and the EM-based post-synaptic sites (left: H01; right: MICrONS) across dendritic branch orders. Each dot corresponds to one branch order.

In the human MTG (H01 dataset), post-synaptic sites were also enriched between the branch orders 3 and 6, yet exhibited a notable extension into higher-order branches compared to mouse (**Figure 4C**). This broader synaptic spread suggests more extensive dendritic integration in human MTG, potentially reflecting increased computational demands of the association cortex. Despite this distal spread, the spatial distribution of post-synaptic sites in the H01 dataset showed a strong and significant correlation with PPSS_PAC_ (**Figure 4D, right**; Pearson’s *r* = 0.89, *p* = 9.59e-22). This high correlation indicates a strong alignment with human synaptic architecture and underscores the robustness of PPSS_PAC_.

Together, these comparisons show that PPSS_PAC_ correlates significantly with synaptic distributions in both mouse and human EM datasets, with particularly strong alignment to human cortical architecture. These results validate PPSS_PAC_ as a biologically grounded model capable of capturing conserved and species-specific features of synaptic organization.

### Morphological subtypes of CA1 cells exhibit distinct and spatially organized synaptic targeting patterns

Previous studies have shown that synapses in hippocampal CA1 neurons are distributed non-randomly across dendritic arbors, with distinct spatial patterns for both excitatory and inhibitory inputs, suggesting a highly organized synaptic architecture^49,50,51^. To better understand the PPSS_PAC_ distributions of CA1 dendrites, we analyzed 2,186 dendrites from the CA1 region and performed hierarchical clustering based on 22 morphological features (**Methods**). This clustering generated two distinct clusters (**Figure 5A**), with optimal statistical separation supported by clustering metrics such as the Silhouette Coefficient (**Methods**). These two clusters, referred to as Cluster 1 and Cluster 2, exhibited different morphological features, suggesting that they represent distinct subtypes of CA1 cells, potentially corresponding to the deep and superficial subpopulations defined along the radial axis^52^.

**Figure 5.**
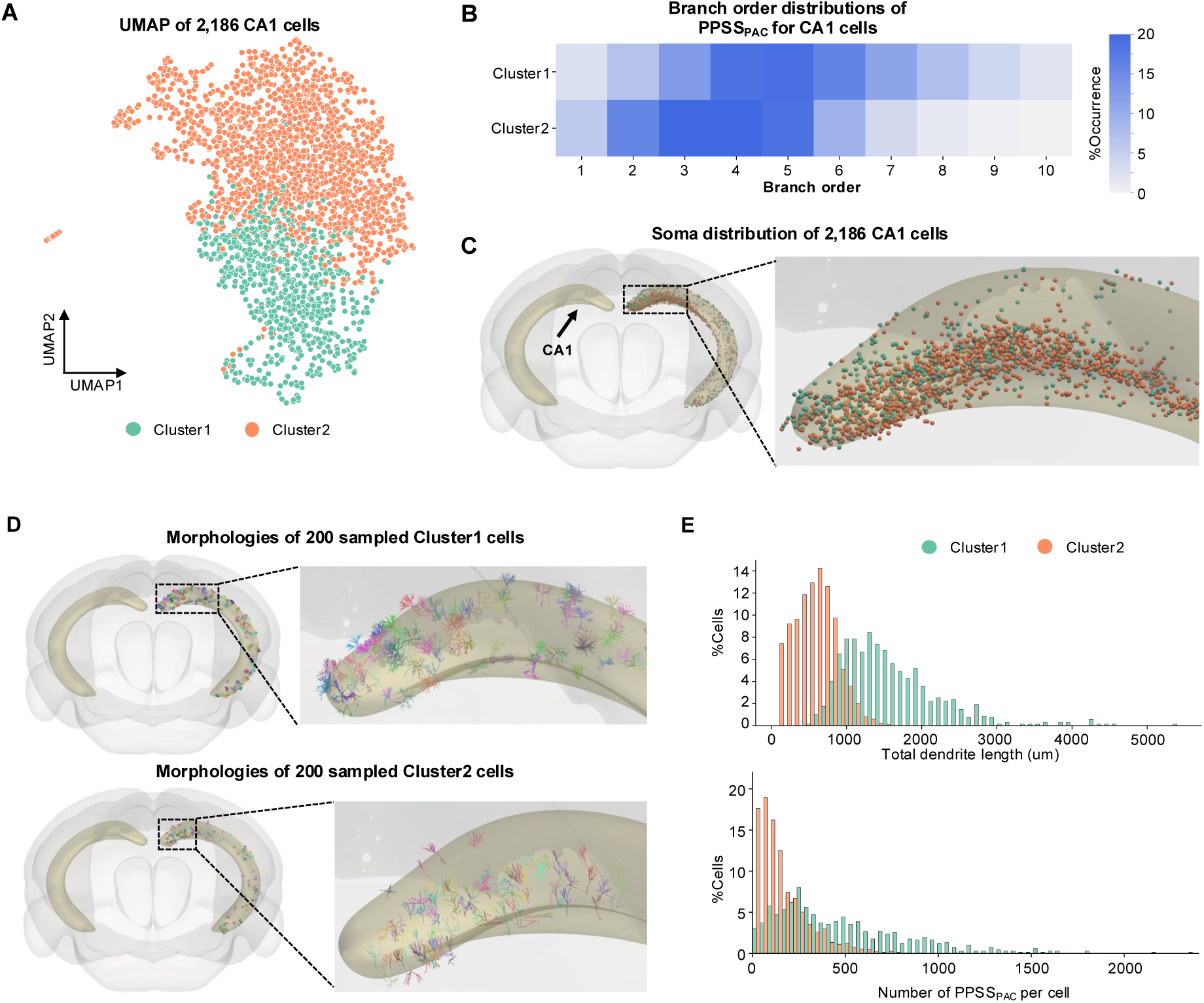
Morphological classification and PPSS_PAC_ distribution of CA1 dendrites. **A.** UMAP embedding of the 2,186 CA1 dendrites. Each dot represents one neuron, colored according to its identified cluster. **B.** The distribution of PPSS_PAC_ across dendritic branch orders for the two CA1 clusters. The colorbar represents the occurrence of PPSS_PAC_ at each branch order. **C.** Spatial distribution of the two CA1 clusters in the mouse brain. Each dot represents the soma location of a CA1 neuron, colored according to its identified cluster. **D.** Example dendritic morphologies of the two CA1 clusters, for the comparison of their morphologies. Each distinct color corresponds to an individual neuron. **E.** The distribution of key morphological characteristics, including total dendritic length (top) and the number of PPSS_PAC_ (bottom) for the two CA1 clusters.

We further examined the distribution of PPSS_PAC_ across dendritic branch orders in the two clusters. The results showed clear differences in how PPSS_PAC_ was distributed across the dendritic branches of the two clusters (**Figure 5B**). PPSS_PAC_ were enriched in distal branches in Cluster1, but were concentrated in proximal branches in Cluster 2. This pattern of post-synaptic targeting suggests that the two dendritic subtypes may receive distinct synaptic inputs, providing a novel descriptor for neuronal classification.

Additionally, we analyzed the spatial distribution of the soma locations for the two clusters within the CA1 region (**Figure 5C**). It turns out that the soma of Cluster 1 was predominantly located in the deeper sub-layer of CA1, closer to the stratum oriens, whereas the soma of Cluster 2 was evenly distributed in the superficial sub-layer, closer to the stratum radiatum. This spatial segregation aligns with a radial axis-dependent organization of CA1 pyramidal cells^53^.

To further characterize the two clusters, we also visualized the dendritic morphologies of the two clusters in the CCFv3^33^. Cluster 1 neurons exhibited longer dendrites compared to Cluster 2 (**Figure 5D**, **E**). Moreover, we compared the total number of PPSS_PAC_ per cell within the two clusters (**Figure 5D**, **E**). Cluster 1 showed a higher number of PPSS_PAC_ per cell than Cluster 2, further validating that the two clusters differ in their synaptic properties. The two features also align closely with the characteristics of deep and superficial pyramidal cells, supporting the hypothesis that these two clusters represent distinct subtypes within the CA1 region^54^. To validate the robustness of our CA1 morphological classification, we performed a control analysis by randomly assigning CA1 cells into two groups. In contrast to the clear spatial segregation observed with morphology-based clustering, these randomly assigned groups showed no significant differences in: (1) the distribution of soma locations across the CA1 region (**Supplementary Figure 6A, B**), (2) the spatial pattern of PPSS_PAC_ (**Supplementary Figure 6C**), or (3) total dendritic length and PPSS_PAC_ number (**Supplementary Figure 6D**). This absence of segregation in the random control confirms that the patterns identified by our clustering method reflect a non-random, biologically meaningful organization.

These findings, including the differences in soma distribution, dendritic morphology, and PPSS_PAC_ spatial pattern, suggest that the two clusters likely represent distinct subtypes of CA1 pyramidal cells, potentially corresponding to deep pyramidal cells and superficial pyramidal cells within the hippocampus^55^.

### PPSS_PAC_ spatial organization, but not synaptic density, correlates with gene expression and whole-brain electrophysiological activity

To assess the biological significance of the PPSS spatial patterning, we performed an integrated cross-modality analysis examining its relationship with molecular and functional profiles (**Figure 6**). We observed that no clear association between inter-regional similarity derived from synapse density and that derived from PPSS_PAC_ patterns (**Figure 6A, left**), despite a strong overall correlation between regional synapse density and PPSS_PAC_ counts (**Figure 3C**). This discrepancy may be attributed to the fact that PPSS_PAC_ counts are unrelated to their spatial patterns (**Figure 6A, middle**).

**Figure 6.**
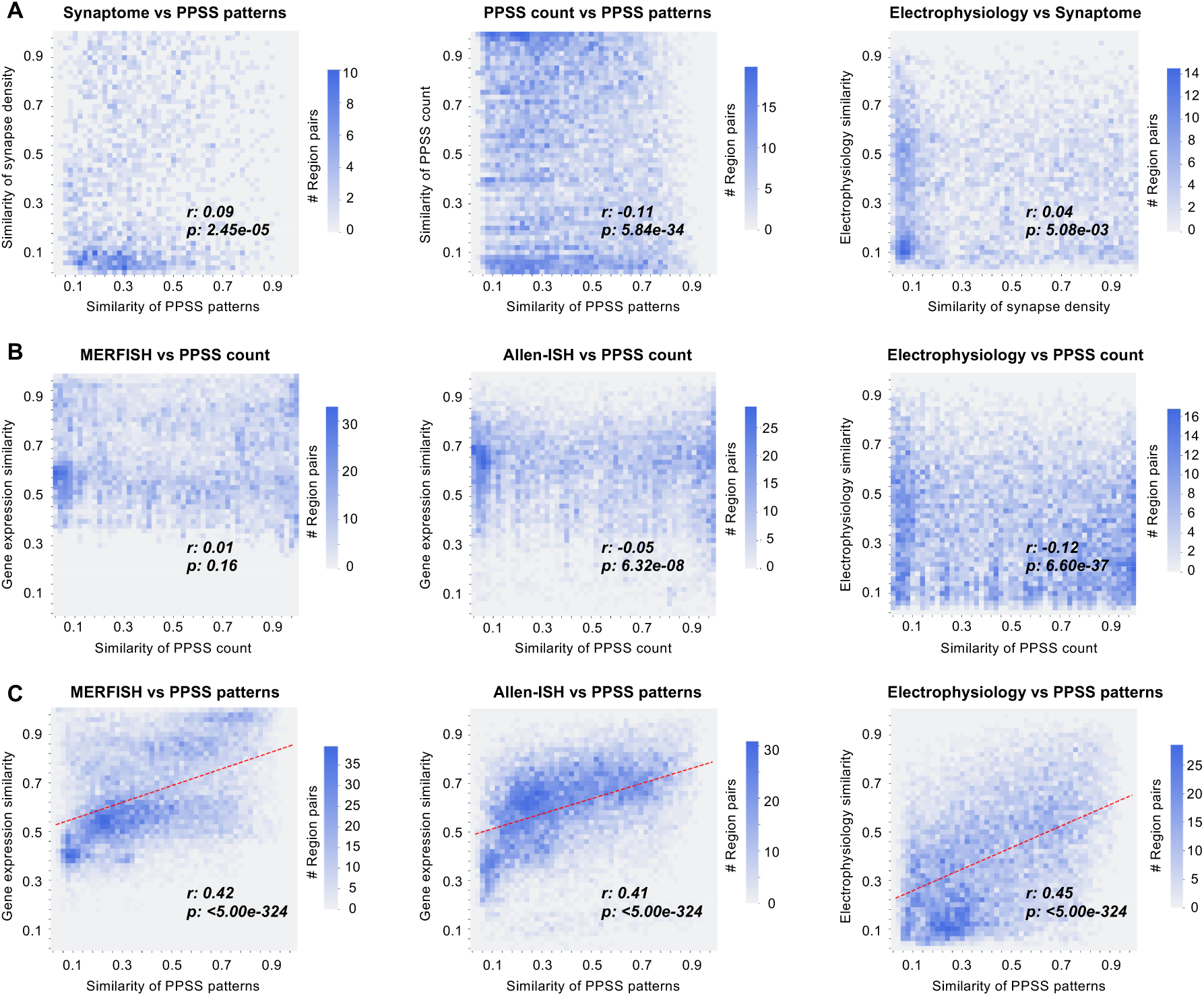
The spatial pattern of the PPSS_PAC_ correlates with gene expression and whole-brain electrophysiological activity. **A.** The inter-regional similarity based on the relative synapse density from synaptome vs. the PPSS_PAC_ patterns (**Left**); regional PPSS_PAC_ counts vs. PPSS_PAC_ patterns (**Middle**); and whole-brain electrophysiology (IBL) vs. relative synapse density synaptome (**Right)**. **B. & C.** The inter-regional similarity based on the MERFISH gene co-expression (**Left**); the Allen-ISH gene co-expression (**Middle**); whole-brain electrophysiology (IBL) (**Right**), correlated against PPSS_PAC_ count (**B**), and the PPSS_PAC_ pattern (**C**). The colorbar represents the numbers of brain region pairs. The red dotted line represents the linear model of best fit from non-linear least squares minimization. Statistical significance for all correlations was assessed using a two-tailed Pearson correlation test without adjustment.

We observed that the regional synapse density was not correlated with whole-brain electrophysiological activity^56^, a direct functional readout of the network, indicating that synaptic density alone is an insufficient for characterizing functional specialization across brain regions (**Figure 6A, right**). Additionally, our analysis revealed no correlation between PPSS_PAC_ count and transcriptomic profiles from both MERFISH^56^ (**Figure 6B, left**) and Allen-ISH^58^ (**Figure 6B, middle**) datasets. Like synapse density, PPSS_PAC_ count exhibited no relationship to whole-brain electrophysiological activity (**Figure 6B, right**), underscoring the independence of this structural metric from global neural dynamics.

Most strikingly, inter-regional similarity in PPSS_PAC_ pattern (**Figure 6C**) showed strong positive correlations with transcriptomic profiles from both MERFISH (**Figure 6C, left**) and Allen-ISH (**Figure 6C, middle**) datasets. This consistent correlation with gene expression data from complementary modalities, spanning from cellular-resolution regional maps to a whole-brain transcriptomic atlas, demonstrates that the spatial organization of PPSS_PAC_ is intrinsically associated with the molecular underpinnings of brain region identity in the mouse. Furthermore, this organizational principle extended to brain function, as the PPSS_PAC_ pattern was also strongly correlated with whole-brain electrophysiological activity (**Figure 6C, right**). Together, these results establish the post-synaptic pattern, not synapse density, as a possible key mediator that bridges molecular signatures, synaptic architecture, and system-level neural computations.

## Discussion

In this study, we developed a statistical framework to predict the locations of post-synaptic sites along dendritic arbors, based on 155,743 reconstructed dendritic morphologies from more than 100 mouse brains. Using this large-scale dataset, we generated more than 17.99 million predicted post-synaptic sites, enabling the profiling of synaptic organization across brain-wide regions at single-neuron resolution. This work addresses a fundamental gap in our understanding of how synaptic inputs are spatially organized along dendrites at the whole-brain scale, which has remained largely unexplored due to the biological complexity of dendritic architectures and the technical limitations of large-scale imaging and reconstruction.

Our analysis reveals conserved patterns of synaptic targeting along dendritic branch orders, which do not only offer consistent observations that proximal and distal dendritic regions often serve distinct functional roles^16,59^, but also identify region-specific variations in post-synaptic density and spatial organization along dendrites, suggesting that local neural circuit and cell-type identity may jointly shape the patterns of post-synaptic inputs. At the same time, our findings indicate that these synaptic patterns can serve as informative descriptors of cell identity, offering complementary features for classification in addition to those based solely on soma location, connectivity patterns involving source-target relationships across brain regions^4^, or local dendritic microenvironments defined by morphology and local context^5^. Even more importantly, we provided the first set of evidences that the spatial patterns, rather than the number, of post-synaptic sites correlates with gene expression and whole-brain electrophysiology. This establishes PPSS spatial organization as a powerful mediator that bridges molecular signatures, synaptic architecture, and system-level neural computations.

Looking ahead, this study can be further improved in several respects. First, our framework could be extended to incorporate predicted synaptic partners^3^, as well as distinguished excitatory and inhibitory post-synaptic sites, as we did previously for a much smaller dataset^1^. Moreover, we could enhance the prediction of the post-synaptic sites using molecular identity, electrophysiological properties, or activity patterns, which could also be produced using antibody conjugation or various imaging experiments. Third, it would be interesting to investigate the dynamic patterns of synaptic organization changing over development, learning, and disease progression. Future work incorporating time-series data might reveal how synapse placement evolves over time. Finally, we envision that these synaptic patterns of many different cell types offer a unique opportunity for hardware implementation of new brain-scale neuromorphic computing algorithms that would perform like “digital twins” of the complex mammalian brains.

In conclusion, we introduce a scalable and generalizable approach to predict brain-wide post-synaptic sites that are associated with several important organization principles. Our findings do not only provide insights into the spatial distribution of post-synaptic sites but also enhance our understanding of how these synaptic patterns are associated with a variety of other neuronal attributes. It suggests that the brain-wide connectivity patterns of neurons go well beyond what current EM or molecular methods alone could offer, presenting possibilities for classifying cell types based on how they are wired, not just where they project to. Future integration with multimodal datasets, including transcriptomics, functional imaging, and developmental atlases, could enhance the interpretability and utility of synaptic mapping, supporting more precise models of circuit computation and plasticity. Ultimately, such efforts could contribute to a more comprehensive theory of neuronal identity and network architecture, offering a crucial foundation for developing digital twin models of the brain to support next-generation artificial intelligence systems.

## Methods

### Single-neuron reconstructions

The 155,743 dendrites and 18,271 axonal reconstructions were retrieved from the NeuroXiv platform^31^. The axonal reconstructions were derived from two separate datasets: the SEU-ALLEN dataset^2,22^ and the ION dataset^30^ (Institute of Neuroscience). The SEU-ALLEN dataset, comprising 1,891 fully reconstructed single neurons, was recently integrated into the NeuroXiv platform. The ION dataset, also ported to the platform, includes reconstructions of 6,357 axonal neurons from the prefrontal cortex and 10,023 axonal neurons from the hippocampus. The dendrites were also generated and cross-validated on the NeuroXiv platform. All neuronal morphologies were registered to the CCFv3 atlas^33^ using the mBrainAligner tool^34^.

### Prediction of post-synaptic sites and their spatial distribution

Predicted postsynaptic sites (PPSS) was generated by evaluating the spatial proximity between the 155,743 dendrites and 18,271 axonal reconstructions. We employed two independent computational approaches to derive PPSS^3^: one based on the potential arbor contact (PPSS_PAC_), and another based on predicted boutons along their axonal arbors (PPSS_PB_). To generate the PPSS_PAC_ dataset, for paired axonal and dendritic arbors, we calculated the shortest Euclidean distance between all axonal compartments and all dendritic compartments. Dendritic compartments within 5 μm of their nearest axonal compartments were identified as PPSS_PAC_. To generate the PPSS_PB_ dataset, we replaced axonal arbors with the putative bouton sites^2,32^ in the above calculations and identified the nearest dendritic compartments within 5 μm of paired bouton sites. The corresponding dendritic compartments were then designated as PPSS_PB_.

To analyze the spatial distribution of these predicted post-synaptic sites, we first determined the order of each dendritic branch based on its bifurcating tree structures. For each neuron, we counted the number of PPSS occurrences at each branch order and converted them into proportions. These proportions were then averaged across all neurons within the same brain region by dividing the sum of proportions at each branch order by the total number of neurons. The averaged proportions across branch orders were used as feature vectors representing the PPSS distribution for each brain region. In addition, we also used an alternative normalization strategy to help correct for the fact that high-order branches are often present in fewer neurons. For each brain region, we counted how many neurons had branches at each order, and then divided the total number of PPSS at each order by that number of neurons.

### The clustering of PPSS_PAC_ distributions for brain regions

To characterize brain regions based on their PPSS_PAC_ distribution across dendritic branch orders (from 1 to 10), each branch order was treated as an individual feature and standardized using the StandardScaler. Subsequently, the UMAP algorithm (umap-learn package, version 0.5.9; function: ’UMAP’; parameters: min_dist=0.5, spread=1.0, metric=’cosine’, n_neighbors=10, random_state=8) was applied to project the standardized PPSS_PAC_ distribution of each brain region into a two-dimensional space. This two-dimensional embedding was used to visualize the similarity relationships among brain regions based on their PPSS_PAC_ distributions.

We used the standardized PPSS_PAC_ distribution features (branch orders 1 to 10) and calculated the Euclidean distance between these features for each pair of brain regions. Pairwise Euclidean distances were then calculated between all region pairs. These distances were converted to inter-regional similarity measures using:

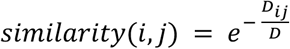

where 𝐷*_ij_* is the Euclidean distance between region *i* and region *j*, and 𝐷 is the median Euclidean distance.

Based on the inter-regional similarity, we performed hierarchical clustering using the scipy package (version 1.13.1; function: ‘linkage’; parameters: method= ‘weighted’, metric= ‘euclidean’) to further analyze the grouping of brain regions according to their PPSS_PAC_ patterns, thereby identifying modules of brain regions with similar PPSS_PAC_ distributions.

### PPSS_PAC_ validations using multi-modality datasets

While the statistical reliability of PPSS_PAC_ has been previously validated^3^, we extended this validation in the current study using multiple complementary approaches and multi-modality datasets.

First, we compared the branch order distributions derived from PPSS_PAC_ and PPSS_PB_. Specifically, we conducted a correlation analysis between the occurrence percentages of PPSS_PAC_ and PPSS_PB_ at corresponding brain regions and branch orders. Although these two methods are based on distinct approaches, we hypothesized a strong positive correlation between them.

Second, we validated PPSS_PAC_ using an independent synaptome dataset^28^ derived from a transgenic mouse model expressing fluorescently labeled postsynaptic proteins PSD95 and SAP102. This dataset includes high-resolution confocal imaging and automated analysis, resulting in a comprehensive, anatomically annotated, brain-wide synapse atlas. For comparison, we selected brain samples from the juvenile to adult stages (P60-P99) and focused on 64 brain regions that overlapped with our PPSS_PAC_ dataset. Synapse densities from this atlas were compared with the corresponding PPSS_PAC_-derived synaptic site counts.

Third, we compared PPSS_PAC_ with our previously generated SEU-net dataset^3^, which provides a map of neuronal connectivity strength across brain regions. Specifically, we examined the relationship between the total signal strength received by a brain region (as the signal recipient) and the number of PPSS_PAC_ sites in that region.

Fourth, we further validated PPSS_PAC_ by comparing it with a brain vascular map^45^. Specifically, we analyzed the relationship between vascular length in each brain region and the number of PPSS_PAC_ sites in that region.

Finally, we validated PPSS_PAC_ against two independent high-resolution EM datasets. The first dataset, from the MICrONS functional connectomics project^27^, includes EM reconstructions of over 200,000 neurons and 0.5 billion synapses in the mouse visual cortex. The second dataset consists of a petascale EM reconstruction of a human temporal cortex sample^8^, which includes thousands of neurons and over 100 million synapses. For both datasets, we compared the distribution of post-synaptic sites on dendritic branches observed in these EM reconstructions with the distribution of PPSS_PAC_ sites, highlighting similarities and differences across species and datasets.

### Clustering of CA1 dendrites

We selected all neurons located in the CA1 region from the 155,743 reconstructed dendrites, resulting in 2,186 CA1 dendrites for morphological analysis. Then we computed 22 morphological features for each dendrite using the ‘global_neuron_feature’ plugin of Vaa3D (version: 3.601). These features included: ‘Nodes’, ‘SomaSurface’, ‘Stems’, ‘Bifurcations’, ‘Branches’, ‘Tips’, ‘OverallWidth’, ‘OverallHeight’, ‘OverallDepth’, ‘AverageDiameter’, ‘Length’, ‘Surface’, ‘Volume’, ‘MaxEuclideanDistance’, ‘MaxPathDistance’, ‘MaxBranchOrder’, ‘AverageContraction’, ‘AverageFragmentation’, ‘AverageParent-daughterRatio’, ‘AverageBifurcationAngleLocal’, ‘AverageBifurcationAngleRemote’, and ‘HausdorffDimension’.

All features were first standardized (z-score normalization), the UMAP algorithm (umap-learn package, version 0.5.9; function: ’UMAP’; parameters: min_dist=2.0, spread=1.0, metric=’ euclidean’, n_neighbors=5, random_state=42) was applied to project the standardized morphological features for each dendrite into a two-dimensional space for visualization.

The PCA algorithm (‘scikit-learn’ package; version: 1.6.1; parameters: n_components=0.99) was used to reduce the dimensionality of the standardized morphological features in the first step. Subsequently, the embeddings were then reduced using Multidimensional Scaling (MDS) for nonlinear projection (‘scikit-learn’ package; version: 1.6.1; parameters: n_components=5, random_state=42). These features were then used for hierarchical clustering using the scipy package (version 1.13.1; function: ‘linkage’; parameters: method= ‘ward’, metric= ‘euclidean’).

To determine the optimal number of clusters, we evaluated clustering results across different cluster numbers using three standard unsupervised metrics: Davies-Bouldin score, Calinski-Harabasz score, and Silhouette score (‘scikit-learn’ package; version: 1.6.1). All three metrics indicated that a two-cluster solution yielded the optimal separation of dendritic types.

### PPSS_PAC_ validation using electrophysiological data

The electrophysiological data was downloaded from the Brain-wide map dataset**^Error!^ ^Reference^ ^source^ ^not^ ^found.^** from the OpenAlyx database using the API of Open Neurophysiological Environment (ONE) python library, which contains the Neuropixels recordings of mouse brain activity acquired during a decision-making task from the International Brain Laboratory (IBL). We used the data that passed the three criteria (median amplitude > 50 uV; noise cut-off < 20 uV; slidingRP_viol = 1) at the cluster level and obtained a total of n=32766 single units in the whole brain. For each single unit, we used firing rate, the minimum, maximum, geometric median of spike amplitudes and the standard deviation of log-amplitudes of spikes for the correlation analysis. Each single unit has also been annotated according to its location.

### Quantification of inter-regional similarity

Inter-regional similarity was computed using complementary metrics tailored to the data type. For gene expression data of MERFISH^56^ and Allen-ISH dataset for each brain region, inter-regional similarity was defined as the Pearson correlation coefficient between the feature vectors between two brain regions.

For features with quantifiable values, such as synapse density or PPSS_PAC_ counts and the electrophysiological data, the inter-regional similarity was quantified using Euclidean distance. Specifically, for each pair of brain regions 𝑖 and 𝑗, we calculated the Euclidean distance 𝐷*_ij_* between their features. To convert this distance into a similarity measure, all pairwise distances were normalized by the median Euclidean distance 𝐷 across the entire matrix.

For correlation analysis, we excluded region pairs when the proportion of near-zero spatial similarity values (< 0.01) exceeded a threshold of 10%, in order to mitigate the influence of extremely skewed distributions.

## Acknowledgment

This project was mainly supported by several grants awarded to H.P., who is a New Cornerstone Investigator and a Shanghai Academy of Natural Sciences (SANS) Senior Investigator. Y.W. was also supported by the National Natural Science Foundation of China (32471148), the Key Research and Development Program of Zhejiang Province (2024C01142), China.

## Author Contribution

H.P. conceptualized and managed this study and instructed the detailed development of methods, experiments, and analyses. Y.W., F.X. and Y.L. generated the analysis results along with biological interpretation. F.L. contributed ideas of analysis of synaptic distributions and their association with EM data, and also implication of bio-realistic synaptic computing using AI hardware. H.P., Y.W., F.X., and Y.L. wrote the initial draft. Y.W. and H.P. reviewed and edited the manuscript with input from all coauthors.

## Declaration of Interests

The authors declare no competing interests.

## Data and Materials Availability

All code, prompt designs, and datasets used in this study are openly accessible at the Zenodo repository (https://zenodo.org/records/18093063) under the MIT License (https://opensource.org/licenses/MIT).

---

**Supplementary Figure 1.**
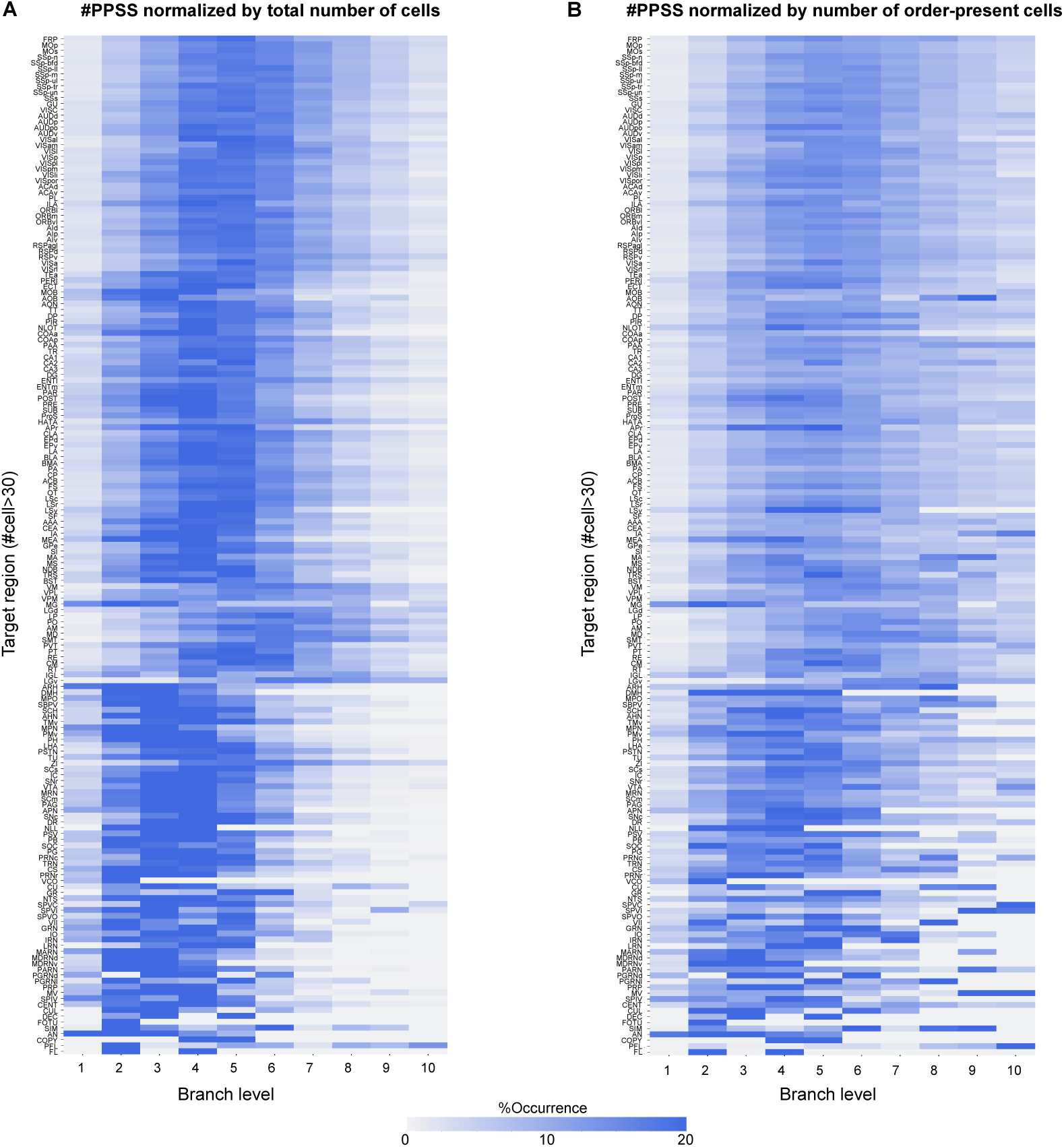
Normalized distributions of PPSS_PAC_ occurrences across dendritic branch orders for 147 brain regions (≥30 cells each). **A.** Heatmap showing the relative proportion of PPSS_PAC_ at each branch order normalized by the total number of neurons sampled from each brain region. **B.** Heatmap showing PPSS_PAC_ proportions normalized by the number of neurons that possessed branches at that specific order. The color bar represents the proportion of PPSS_PAC_ at each dendritic branch order.

**Supplementary Figure 2.**
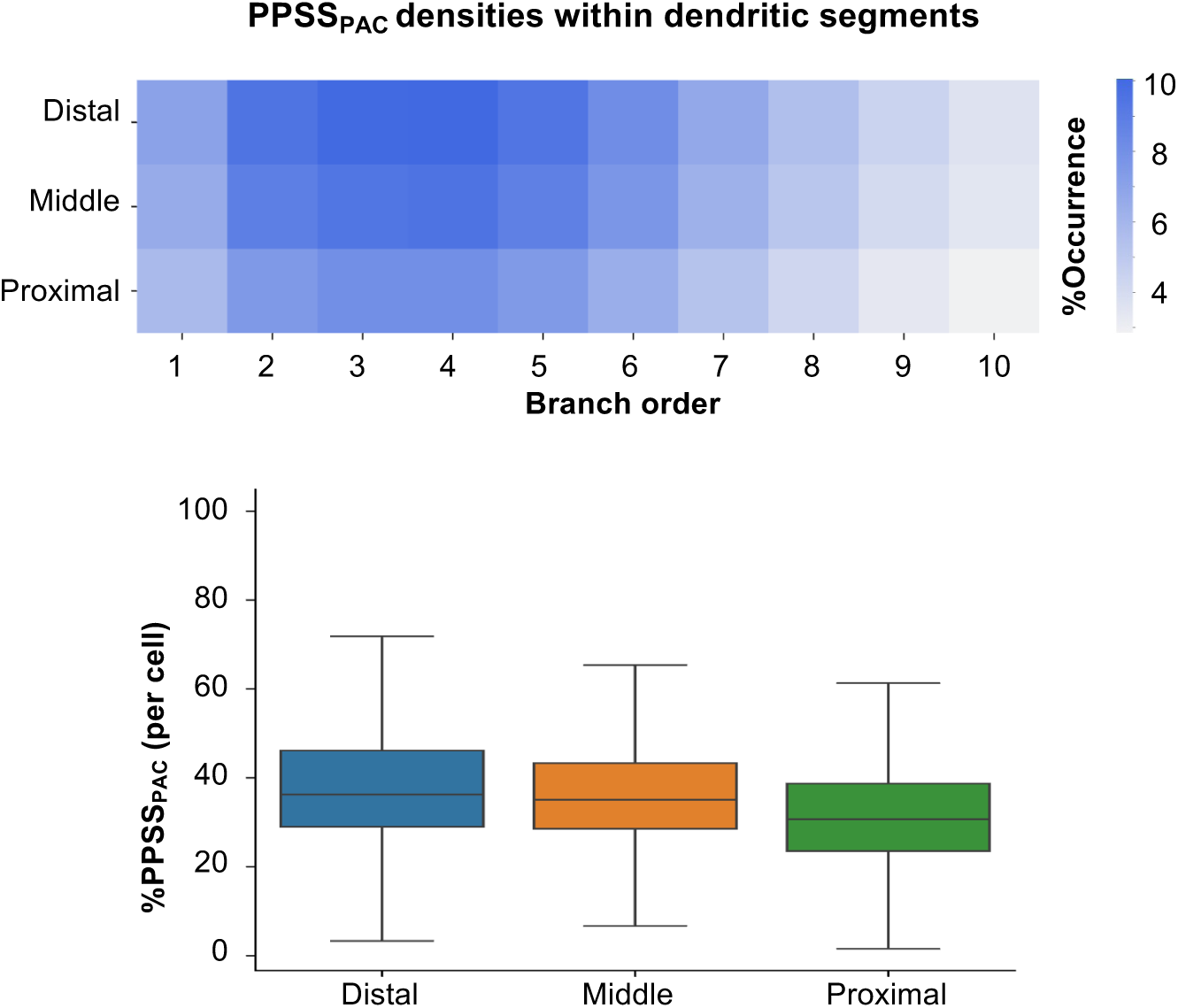
The proportion of PPSS_PAC_ across three dendritic segments: proximal (near the branch origin), middle (near the branch center), and distal (near the branch end). Box plots show the center line as the median, and box limits as upper (75%) and lower (25%) quartiles.

**Supplementary Figure 3.**
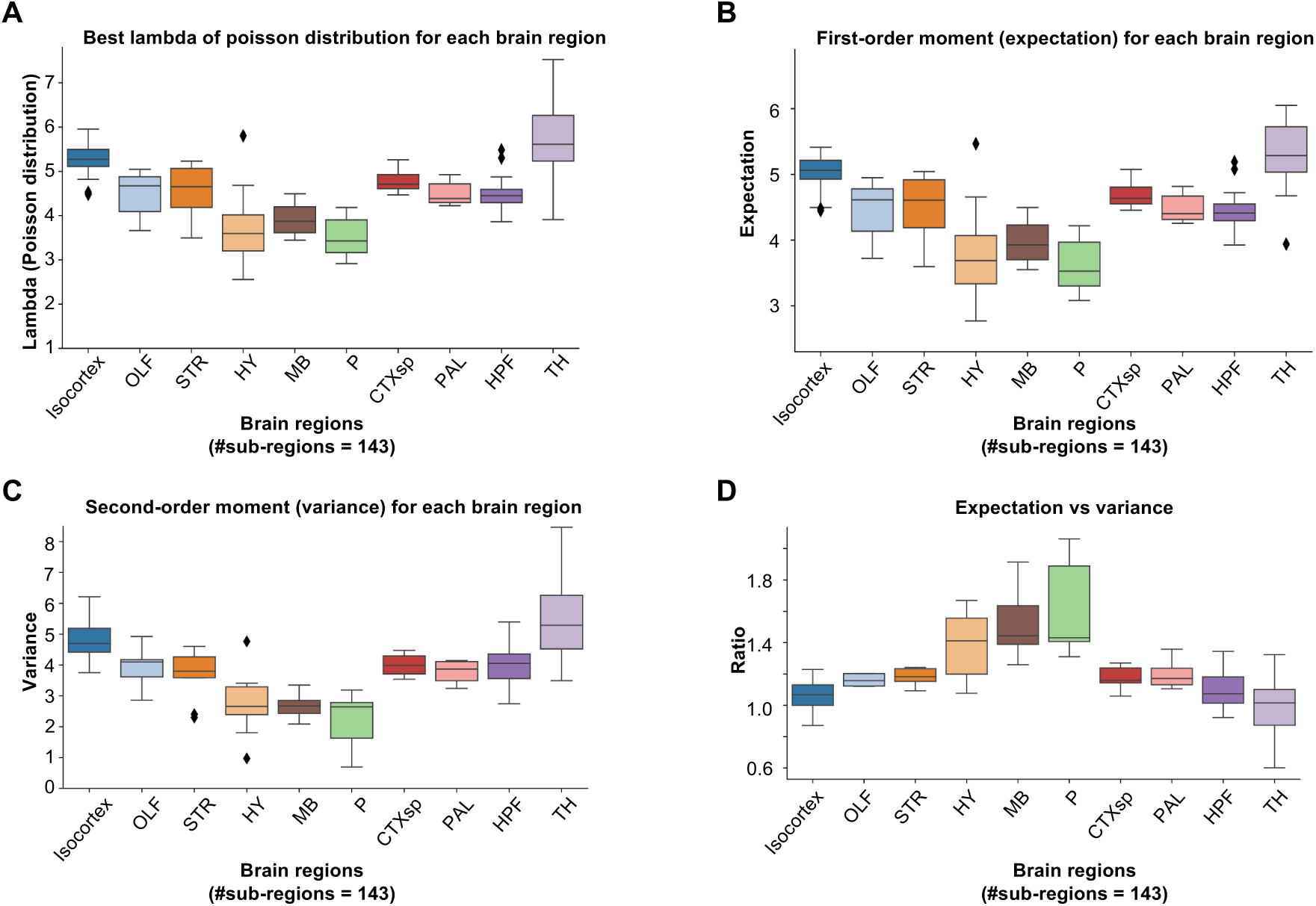
Parameters of the Poisson models fitting the PPSS_PAC_ distributions and related statistics. **A.** Distributions of the estimated λ parameters of the best-fitting Poisson models for sub-regions within each major brain region. **B.** Distributions of the expected values (means) of the Poisson fits for sub-regions within each major brain region. **C.** Distributions of the variances of the Poisson fits for sub-regions within each major brain region. **D.** Distributions of the ratios between the expected values and variances of the Poisson fits for sub-regions within each major brain region.

**Supplementary Figure 4.**
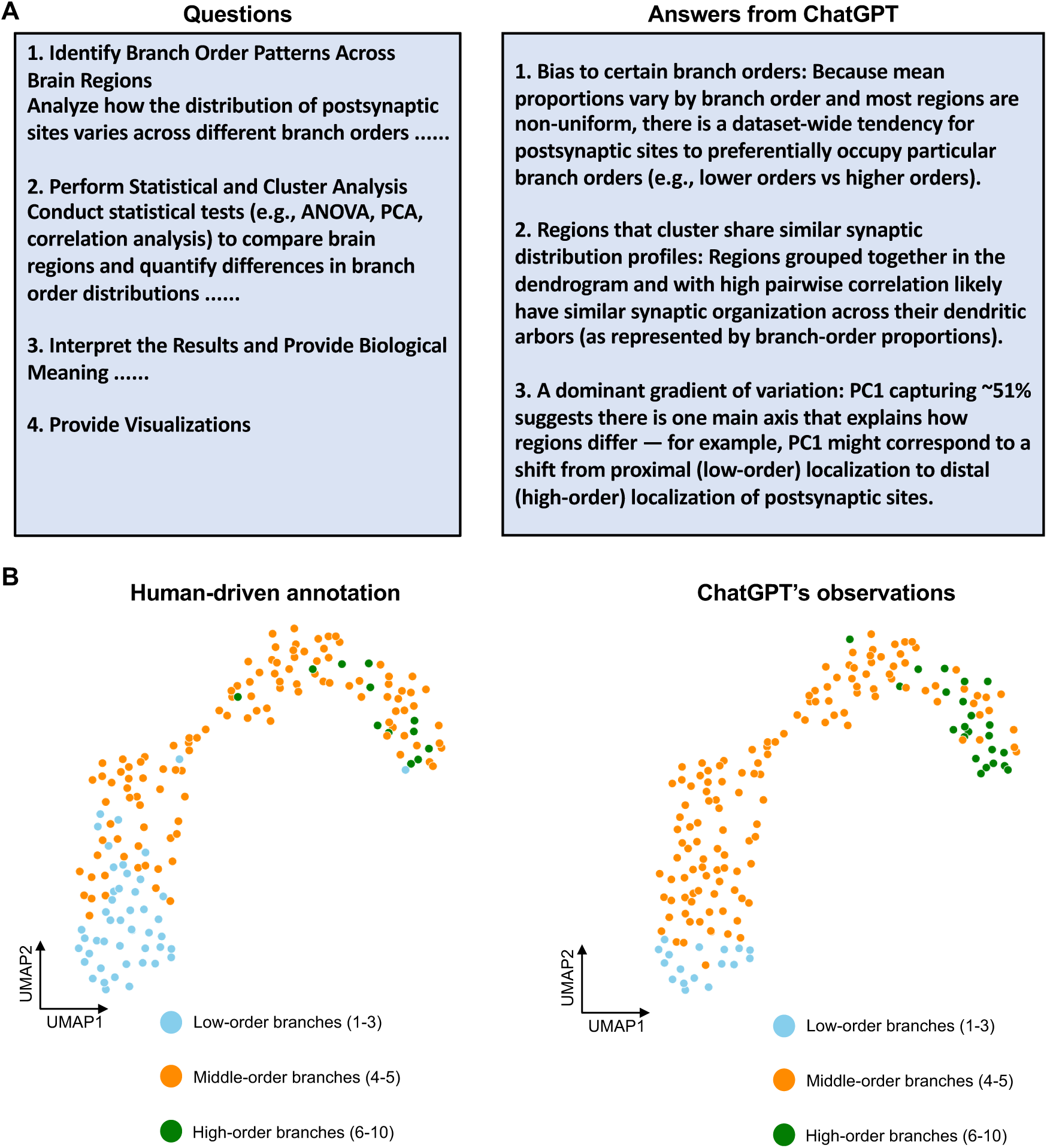
Using ChatGPT to investigate the branch order distributions of PPSS_PAC_ **A**. Question design and answers from ChatGPT. **B**. UMAP layout of 174 brain regions using their branch order distributions of PPSS_PAC_. Left: brain regions are categorized by human-driven annotation; Right: brain regions are categorized by ChatGPT’s observations. Each dot represents a brain region, colored according to the branch order at which its peak occurrence is observed.

**Supplementary Figure 5.**
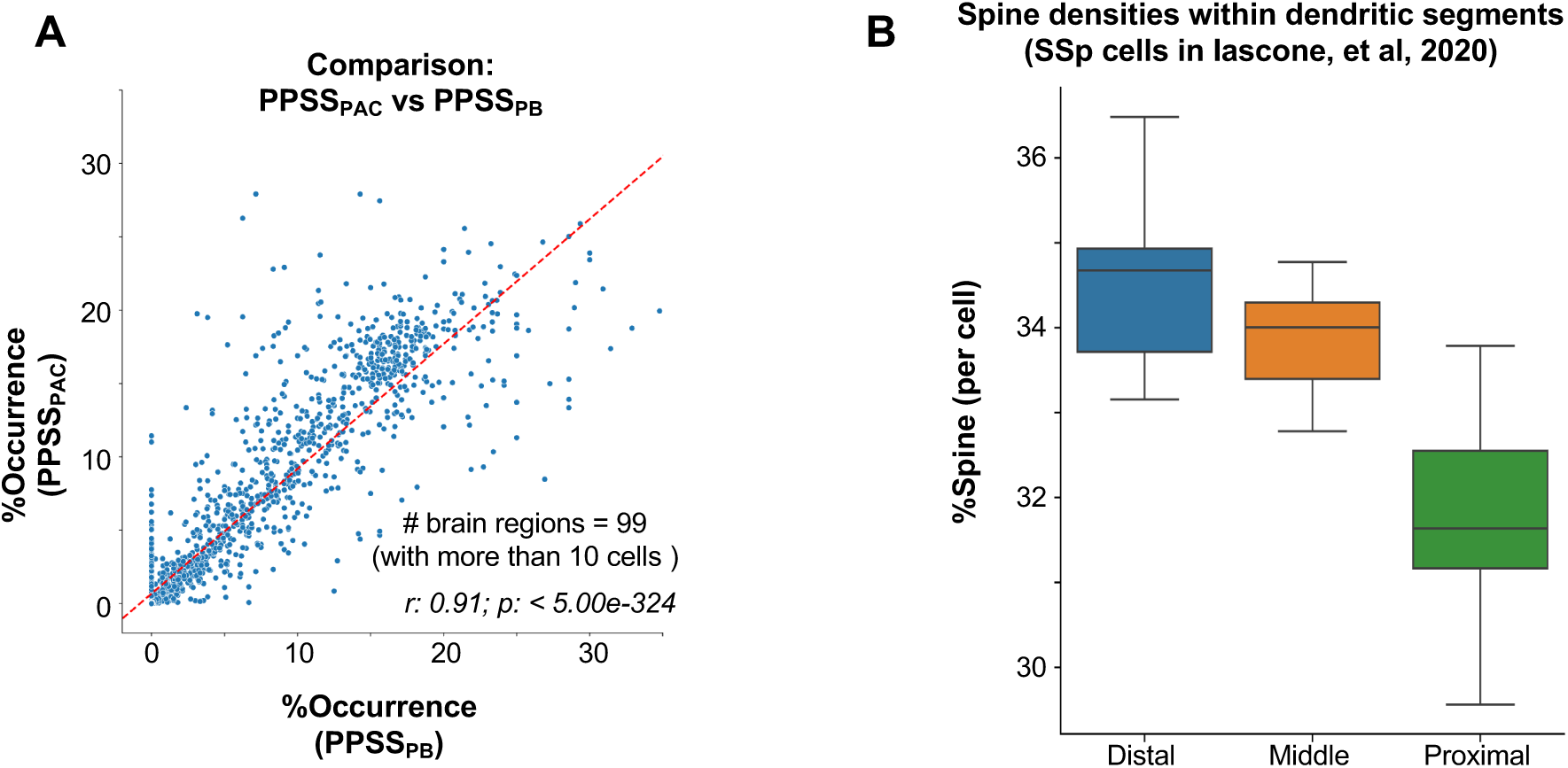
Cross-validation of PPSS_PAC_ using multiple datasets. **A.** Correlation of anatomical distributions between PPSS_PAC_ and PPSS_PB_. **B.** Spine densities within dendritic segments for SSp neurons.

**Supplementary Figure 6.**
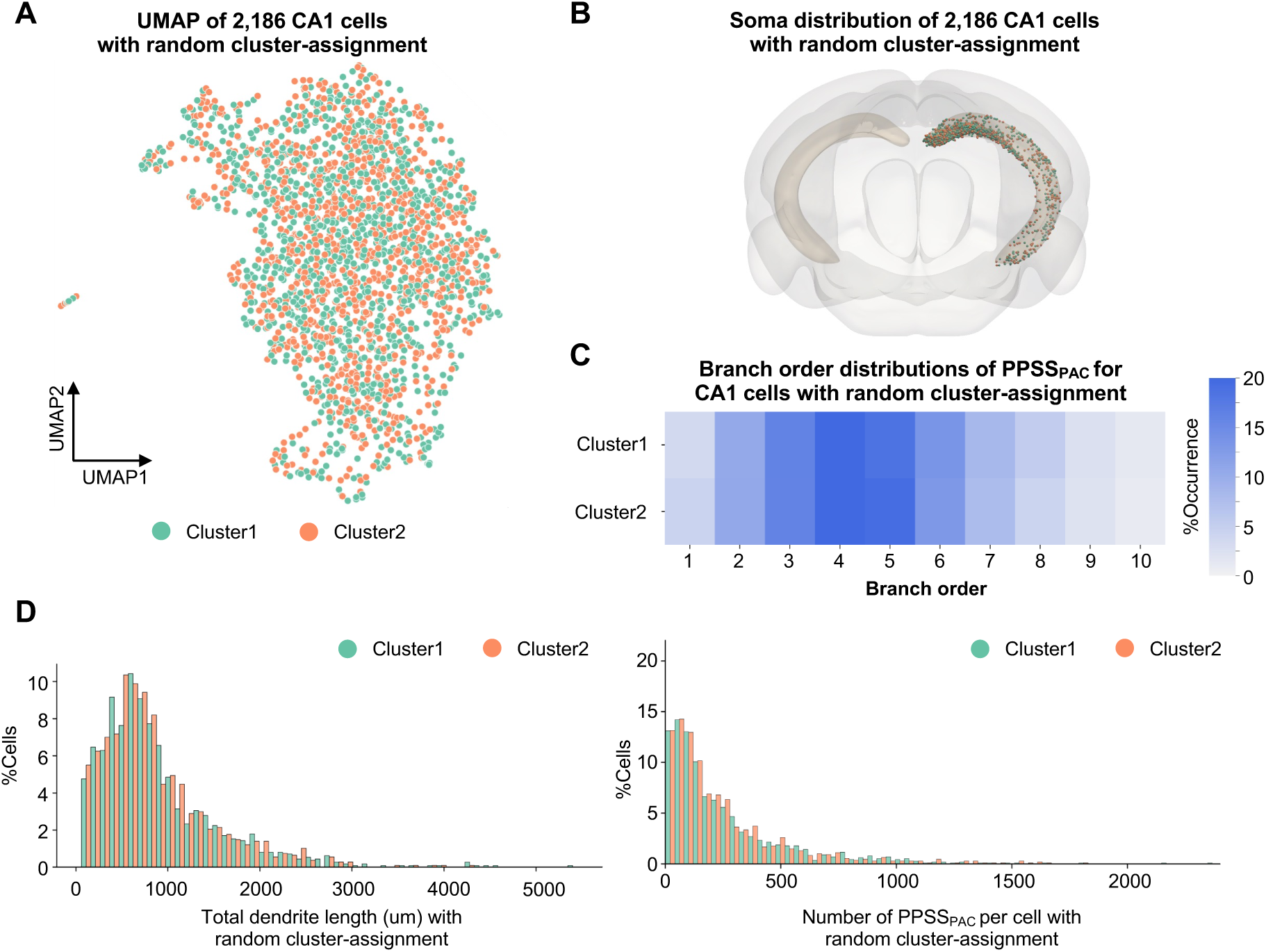
Control analysis by randomly separating 2,186 CA1 cells into two groups. **A.** UMAP projection of 2,186 CA1 neurons, randomly divided into two equal-sized groups (Cluster 1 and Cluster 2). Each point represents a single neuron in the reduced 2D space. **B.** The soma locations of two groups of CA1 cells in the CCF brain. **C.** Spatial distribution of the two groups in the mouse brain. **D.** The distribution of key morphological characteristics, including total dendritic length (left) and the number of PPSS_PAC_ (right) for the two groups.

## Notes

### Competing Interest Statement

The authors have declared no competing interest.

